# *midline* represses Dpp signaling and target gene expression in *Drosophila* ventral leg development

**DOI:** 10.1101/2021.12.22.473919

**Authors:** Lindsay A. Phillips, Markle L. Atienza, Jae-Ryeon Ryu, Pia C. Svendsen, Lynn K. Kelemen, William J. Brook

## Abstract

Ventral leg patterning in *Drosophila* is controlled by the expression of the redundant T-box Transcription factors *midline (mid)* and *H15*. Here we show that *mid* represses the Dpp-activated gene *Daughters against decapentaplegic* (*Dad*) through a consensus TBE site in the minimal enhancer, Dad13. Mutating the Dad13 DNA sequence results in an increased and broadening of Dad expression. We further demonstrate that the engrailed-homology-1 domain of Mid is critical for regulating the levels of phospho-Mad, a transducer of Dpp-signaling. However, we find that *mid* does not affect all Dpp-target genes as we demonstrate that *brinker* (*brk*) expression is unresponsive to *mid*. This study further illuminates the interplay between mechanisms involved in determination of cellular fate and the varied roles of *mid*.

**Summary statement:** Ventral patterning is controlled in part by the T-box Transcription factor *midline* blocking Dpp signaling and Dpp-activated genes, though *midline* does not affect the Dpp-repressed gene *brk*.

## Introduction

Proper organization of tissue is crucial for maintaining the body plan of animals. This organization occurs during development when multiple factors cooperate to determine cell fate. In particular, the *Drosophila melanogaster* leg is organized in part by the interplay between signaling molecules and transcription factors. Dorsal and ventral fates in *Drosophila* legs are dependent on the action of morphogens and selector genes. The morphogens *decapentaplegic* (*dpp* – a Bone Morphogenetic Protein (BMP) homolog) and *wingless* (*wg* – a fly Wnt) are induced by *Hedgehog*-signaling. *Wg* is induced in the ventral domain and controls ventral fate through induction of the redundant Tbx20 class T-box transcription factor homologs *midline (mid)* and *H15*, that specify ventral fate (Svendsen et al., 2009). Dorsal fate is dictated by *dpp*, which is expressed at high levels in the dorsal domain and low levels in the ventral domain, though it has no role in ventral fate aside from joint formation (Held and Heup, 1996; Manjón et al., 2007). Dpp-signaling is mediated by transcription factors *Mothers against dpp* (*Mad*), a fly Smad1/5 activator and *Medea* (*Med*), a fly Co-Smad (Hudson et al., 1998; Kim et al., 1997).

*mid* and *H15* control ventral patterning being both necessary and sufficient to specify the fate in the ventral region of fly legs (Svendsen et al., 2009). They act as selector genes, transcription factors that specify cell fate for a particular developmental region, with expression restricted to the region in which they specify cell fate (Crickmore and Mann, 2008). The ventral specific expression of *mid* and *H15* is controlled through a combination of Wg activation and Dpp repression (Svendsen et al., 2009; Svendsen et al., 2015). When *mid* and *H15* function is lost in the ventral leg, tissues are transformed into dorsal while ectopic expression of *mid* or *H15* induces ventral fate in dorsal regions (Svendsen et al., 2009).

How does *mid* control ventral development? We have shown that one role of *mid* is to block Dpp signaling in the ventral domain (Svendsen et al., 2019). The distribution of phosphorylated Mad (pMad) follows the same pattern as *dpp*, with significant levels of staining in the dorsal domain and weaker staining in ventral region, indicating that the Dpp pathway is activated at lower levels in ventral cells. While Dpp does not contribute to ventral patterning, double mutant analysis shows *mid H15* mutant clones blocked for Dpp signaling are rescued in some regions of the leg, indicating an inhibitory effect of *mid* on Dpp signaling that is necessary for ventral patterning. This is further demonstrated by the inhibition of pMad accumulation by *mid* (Svendsen et al., 2019). Additionally, *mid-*expressing clones repress the Dpp-target genes *Dad, Upd3*, and *mid* itself in an eh1-dependent manner, and using chromatin-immunoprecipitation (ChIP) assays, Mid has been shown to localize to enhancers for these genes (Svendsen et al., 2019).

Here, we study further the way *mid* antagonizes Dpp signaling, showing that it represses the regulation of *Dad*, a target gene activated by Dpp/pMad, but has no effect on *brk*, a gene repressed by Dpp/pMad. We show that Mid repression of *Dad* depends on a predicted T-box binding element (TBE) in the Dad13 enhancer, and that Mid-inhibition of Dpp-dependent pMad accumulation and tissue re-patterning depends on the eh1 repressor domain.

## Results

### Mid blocks Dpp activation of Dad13 reporter expression

Our previous work showed that the eh1 repressor binding domain was required for Mid selector gene function. We also showed that Mid acted by antagonizing Dpp signaling, reducing the levels of pMad accumulation in the ventral domain of the fly leg (Svendsen et al., 2019). Here we further investigate how *mid* affects Dpp-target gene expression. The Dpp-target genes *Dad* and *brk* are canonical examples of genes activated or repressed by Dpp signaling, respectively (Gao et al., 2005; Tsuneizumi et al., 1997). Since *mid* antagonizes Dpp signaling, we sought to understand how *mid* regulates both genes. We showed previously that an enhancer-trap reporter of *Dad* (*Dad*-lacZ) was weakly repressed by Mid in an eh1-dependent manner (Svendsen et al., 2019). Mid likely antagonizes Dpp-target genes through direct repression because Mid has been shown to bind to several enhancer fragments of *Dad* in ChIP assays (Svendsen et al., 2019). One of these fragments was a well-characterized 520 bp enhancer, Dad13. The Dad13 expression pattern is similar to that of *Dad*-lacZ, with strong expression in the dorsal domain and weaker expression in the ventral domain. However, the ventral expression of Dad13 is weaker compared to wildtype *Dad* (Fig. S1). We first confirmed that a Dad13 reporter was regulated by Mid. We investigated if loss of *H15* and *mid* function would affect Dad13 expression by generating *H15 mid* loss-of-function clones via mitotic recombination in imaginal discs of second instar larvae. Dad13 expression was detected by RFP expression while clones null for *H15 mid* were marked by the absence of GFP. Ventrally located *H15 mid* loss-of-function clones showed either an increase of Dad13 expression or an expansion into the lateral region of the leg imaginal disc (Fig. 1A,B). Because the *mid* expression domain completely encompasses the ventral Dad13 domain, we induced ectopic Dpp-signaling in lateral regions of the imaginal disc in order to induce Dad13 expression outside the *mid-*expression domain. We then introduced *mid* expression to assess the ability of Mid to repress Dad13. By using AyGAL4, a construct that generates random clones expressing Gal4 under the control of the *actin5C* promoter, we induced clones marked by GFP expression that also expressed a constitutively active form of the Dpp receptor, *thickveins* (UAS-*tkv^QD^*) and/or expressed Mid. As expected, UAS-*tkv^QD^* expressing clones, which are constitutively activated for Dpp-signaling, had increased Dad13-nRFP expression. However, co-expressing UAS-*mid*^+^ and UAS-*tkv^QD^* blocked the ectopic activation of Dad13-nRFP in clones located throughout the disc (Fig. 1C,D). This indicates that the presence of Mid blocks the effects of Dpp-signaling on Dad13 expression. Together these results confirm that Mid does indeed regulate the Dpp-target gene *Dad* via its enhancer fragment, Dad13.

**Figure 1.**
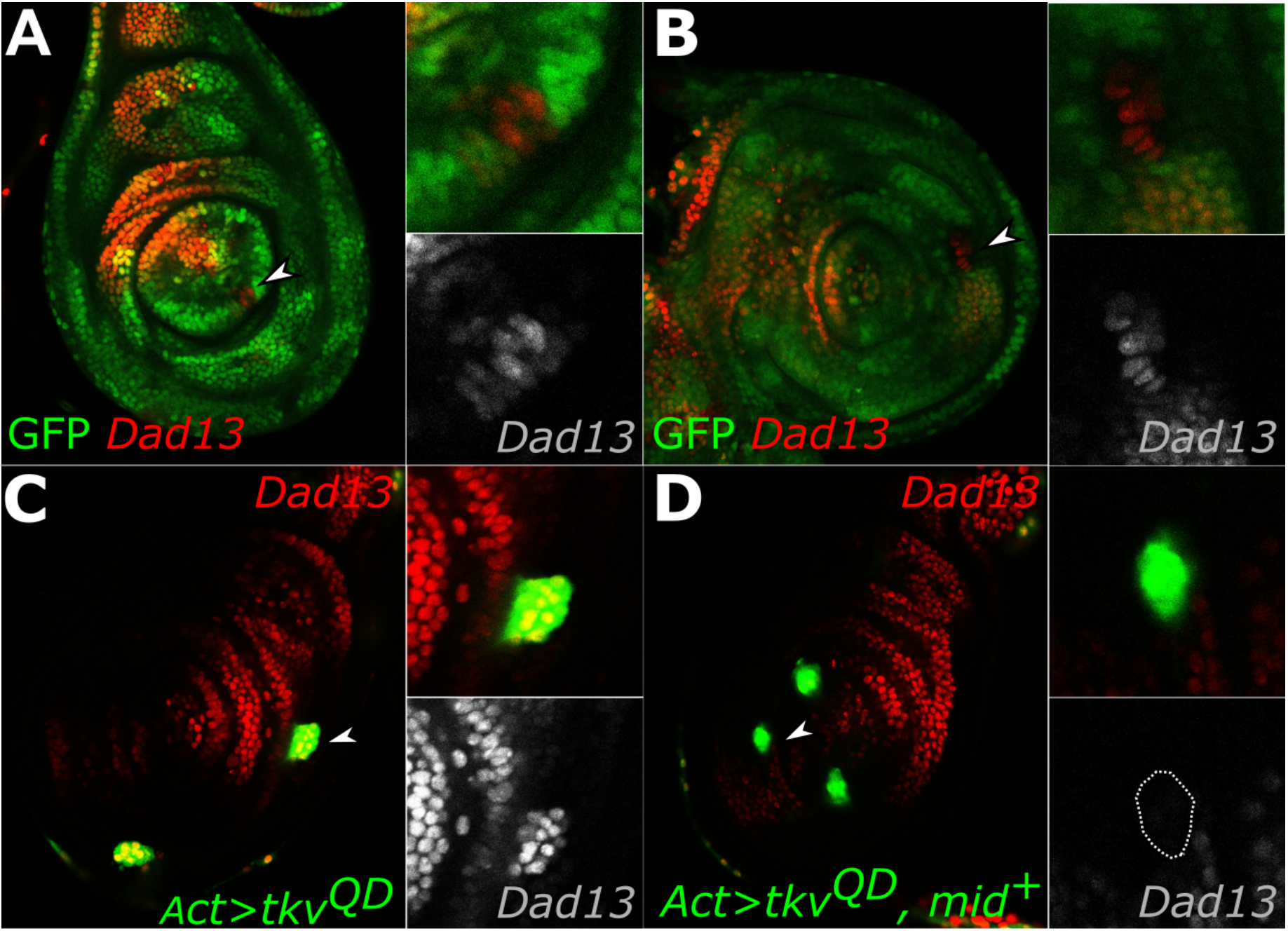
Mid blocks Dpp activation of Dad13 expression. (A-B) Third-instar legs discs expressing Dad13-nRFP (red, single channel) and loss-of-function *H15* and *mid* (lack of green) clones. Clones lacking *H15* and *mid* (A) have increased Dad13-nRFP expression (white arrowhead, lower inset), or (B) expanded Dad13-nRFP expression (white arrowhead, lower inset). (C) Discs with UAS-*tkv^QD^* gain-of-function clones (green) driven by AyGal4 driver showed ectopic Dad13 expression (red, lower inset), while (D) clones co-expressing UAS-*tkv^QD^* and UAS-*mid*^+^ (green) did not induce ectopic Dad13 expression (red, lower inset clone outline).

### *The TBE in Dad13 is necessary for* mid *repression of* Dad

To further investigate the role of *mid* in *Dad* regulation, we examined how Mid influences Dad13 expression. The Dad13 enhancer contains multiple binding sequences for Mad binding elements that are responsible for driving Dad13 expression (Weiss et al., 2010). Dad13 also contains tandem Smad binding element (SBE) sequences (GTCTGTCT) that have a minor role in activating Dad13 in embryos but which have not been tested in leg imaginal discs (Weiss et al., 2010). Adjacent to the SBE sites and separated by a single nucleotide is a consensus T-box binding element (TBE) (AGGTGA) similar to the consensus binding site for Mid (Najand et al., 2012). The proximity of activating SBE next to the potential Mid binding element suggests the activation of Dad13 by Dpp may be blocked by Mid. To test whether the TBE is necessary for Mid regulation of Dad13, we mutated the Dad13 TBE site (Dad13^TBE^) and tested expression with a lacZ reporter (Fig. 2A). Dad13^TBE^ expression was stronger and broader in the ventral domain compared to the Dad13 control (Fig. 2B,D, Fig. S4). This clearly indicates that the TBE is required for repression of *Dad*. Furthermore, the expression of Dad13^TBE^ in dorsal regions outside the *mid* expression domain was more intense compared to Dad13, suggesting that factors other than Mid may also regulate *Dad* through the TBE site. Mutating the SBE (Dad13^SBE^) had no observable effect on Dad13 expression and mutating both the TBE and SBE (Dad13^SBE+TBE^) produced similar expression effects to mutating the TBE alone (Fig. S3). Thus, the SBE site does not appreciably affect *Dad* expression in the leg imaginal disc. To test whether Mid could still regulate expression of Dad13^TBE^ construct, we generated UAS-*mid*^+^ gain-of-function clones and measured Dad13^TBE^ lacZ expression. Consistent with our previous work (Svendsen et al., 2019), *mid* partially repressed the expression of Dad13 in most *mid*-expressing clones in the ventral domain (7/8) (Fig. 2C,C’). In contrast, few *mid*-expressing clones in the ventral domain of discs repressed Dad13^TBE^ (1/11 clones) (Fig. 2E,E’). These results indicate that the wild type TBE in the Dad13 enhancer fragment is an essential element for the *mid*-mediated repression of *Dad*.

**Figure 2.**
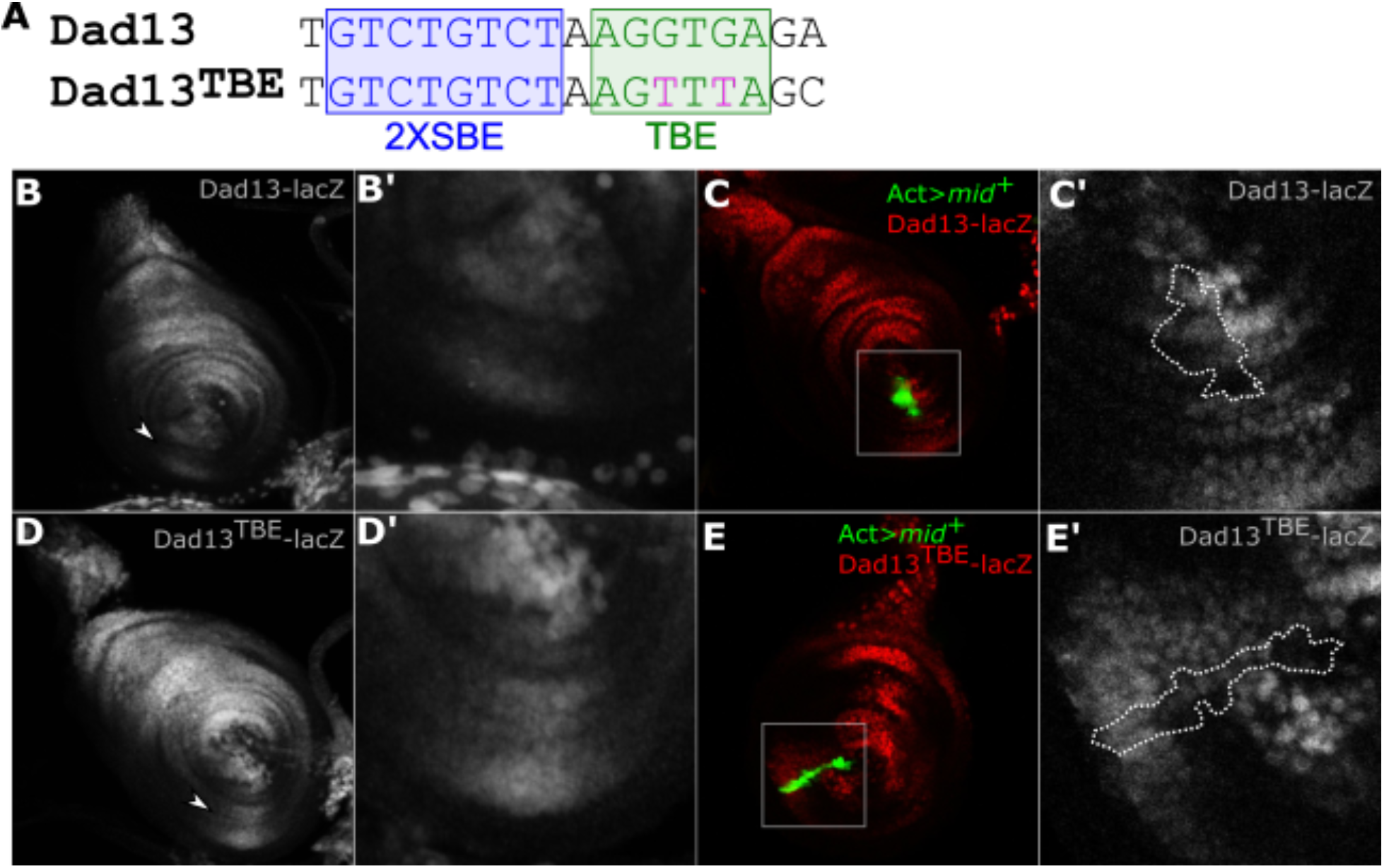
Mid repressed Dad in part though the TBE. (A) Dad13 enhancer fragment sequences, showing the 2X SBE sequence (blue) and the TBE sequence (green). Two G to T substitutions in the TBE were generated for the Dad13^TBE^ construct (fuchsia). (B) Dad13 expression was detected by lacZ with strong staining in the dorsal domain and weak staining in the ventral domain (arrowhead). (B’) Magnified image of ventral expression of Dad13-lacZ. (C, C’) UAS-*mid*^+^ gain-of-function clones (GFP) partially repressed Dad13 expression (red channel, outline, C’). (D) Expression of Dad13^TBE^ is broader and more intense in the ventral domain relative to Dad13 (arrowhead). Dorsal expression is also stronger in Dad13^TBE^ compared to Dad13. (D’) Magnified image of ventral expression of Dad13^TBE^-lacZ. (E, E’) Example of a UAS-*mid*^+^ gain-of-function clone (GFP) which maintains normal levels of ventral Dad13^TBE^ expression. No repression of Dad13^TBE^-lacZ was seen compared to adjacent regions outside the clone (outline, E’)

### mid *does not affect Dpp repression of* brk

*brk* is negatively regulated by Dpp signaling. Unlike *Dad*, which is activated by a complex of pMad and Med, *brk* is repressed by a complex of pMad and Med binding along with a third protein, Schnurri, through a repressive binding element (Hamaratoglu et al., 2014; Marty et al., 2000; Saller and Bienz, 2001; Weiss et al., 2010). In leg imaginal discs, high levels of *brk* and pMad expression are reciprocal, demonstrating their antagonistic activities (Müller et al., 2003) (Fig. S1). We found that neither *H15 mid* loss-of-function clones nor *mid*^+^-expressing gain-of-function clones had any effect on *brk* expression. In loss-of-function experiments, endogenous *brk* expression was detected by a *brk* antibody and remained unchanged in clones lacking *H15 mid* (Fig. 3A, insets). Moreover, *mid*^+^ gain-of-function clones marked by GFP (UAS-*mid*^+^) did not affect *brk* expression as detected by lacZ (Fig. 3B). Because Dpp-signaling represses *brk* while *mid* represses *Dpp*-signaling, we tested if ectopic expression of Dpp signaling or *mid* would increase or otherwise affect *brk* expression. As expected, gain-of-function clones expressing UAS-*tkv^QD^* gain-of-function completely repressed *brk* expression (Fig. 3C). However, clones co-expressing UAS-*mid*^+^ with UAS-*tkv^QD^* had no influence on the *tkv* gain-of-function effect on *brk* expression, which remained completely repressed (Fig. 3D). We have previously demonstrated that clones co-expressing UAS-*mid*^+^ and UAS-*tkv^QD^* markedly reduce pMad staining compared to clones expressing UAS-*tkv^QD^* alone (Fig. S2) (Svendsen et al., 2019), suggesting that there is sufficient residual pMad activation to repress *brk*. These results indicate that *mid* does not regulate *brk* and therefore *mid* does not affect all Dpp-target genes.

**Figure 3.**
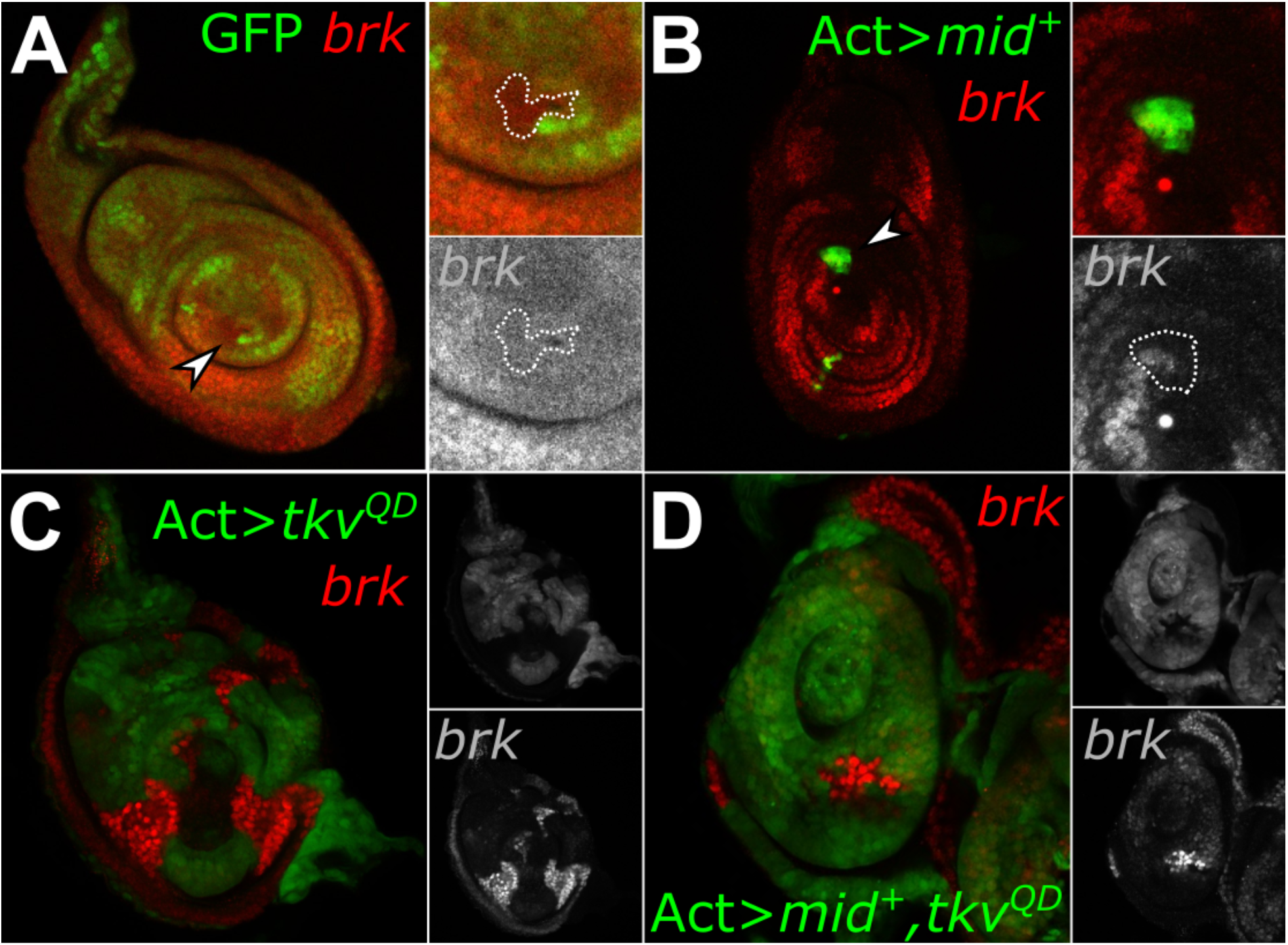
Mid does not affect *brk* function. (A) *H15 mid* loss-of-function (lack of green, arrowhead) clones have no effect on *brk* expression, as detected by a *brk* antibody (red, lower inset clone outline). (B) Discs expressing AyGal4 UAS-*mid*^+^ gain-of-function clones (green) do not affect *brk*-lacZ expression, as detected by anti-β-Galactosidase staining (red, lower inset clone outline). (C-D) Clones expressing UAS-*tkv^QD^* (green, top inset) have a dramatic repressive effect on *brk*-lacZ (red, lower inset), while clones co-expressing UAS-*tkv^QD^* and UAS-*mid*^+^ (green, top inset) did not affect *brk*-lacZ expression (red, lower inset).

### Mid antagonizes Dpp signaling in an eh1-dependent manner

Mid acts as a repressor through its engrailed-homology-1 (eh1) domain, which recruits the co-repressor *groucho (gro)* (Formaz-Preston et al., 2012). The eh1 domain is essential for Mid-mediated repression of genes in the ventral leg including the Dpp-target gene *Dad* (Svendsen et al., 2019). Here we asked if the eh1 is also necessary to inhibit the effects of ectopic Dpp signaling, including phenotypic defects and pMad accumulation. We generated gain-of-function UAS-*tkv^QD^* clones in developing imaginal discs in second instar larvae. In adult legs, these clones gave rise to rounded outgrowths characteristic of ectopic Dpp-signaling (Fig. 4A) (Svendsen et al., 2019). Next, we co-expressed UAS-*tkv^QD^* and a Flag-tagged *mid* (UAS-*mid*^+^-Flag) in adult legs. These clones had fewer defects, which were less severe and had generally smaller outgrowths (Fig. 4C). However, when we co-expressed UAS-*tkv^QD^* with UAS-*mid^eh1^*-Flag, a transgene in which the eh1 domain is mutated and has reduced Gro-binding (Formaz-Preston et al., 2012), the adult legs (Fig. 3E) displayed outgrowths similar in size and severity to clones expressing UAS-*tkv^QD^* alone (Fig. 4A). Overall, more defects were detected in legs expressing either UAS-*tkv^QD^* or UAS-*tkv^QD^* and UAS-*mid^eh1^*-Flag clones as compared to legs expressing UAS-*tkv^QD^* and UAS-*mid*^+^-Flag clones.

**Figure 4.**
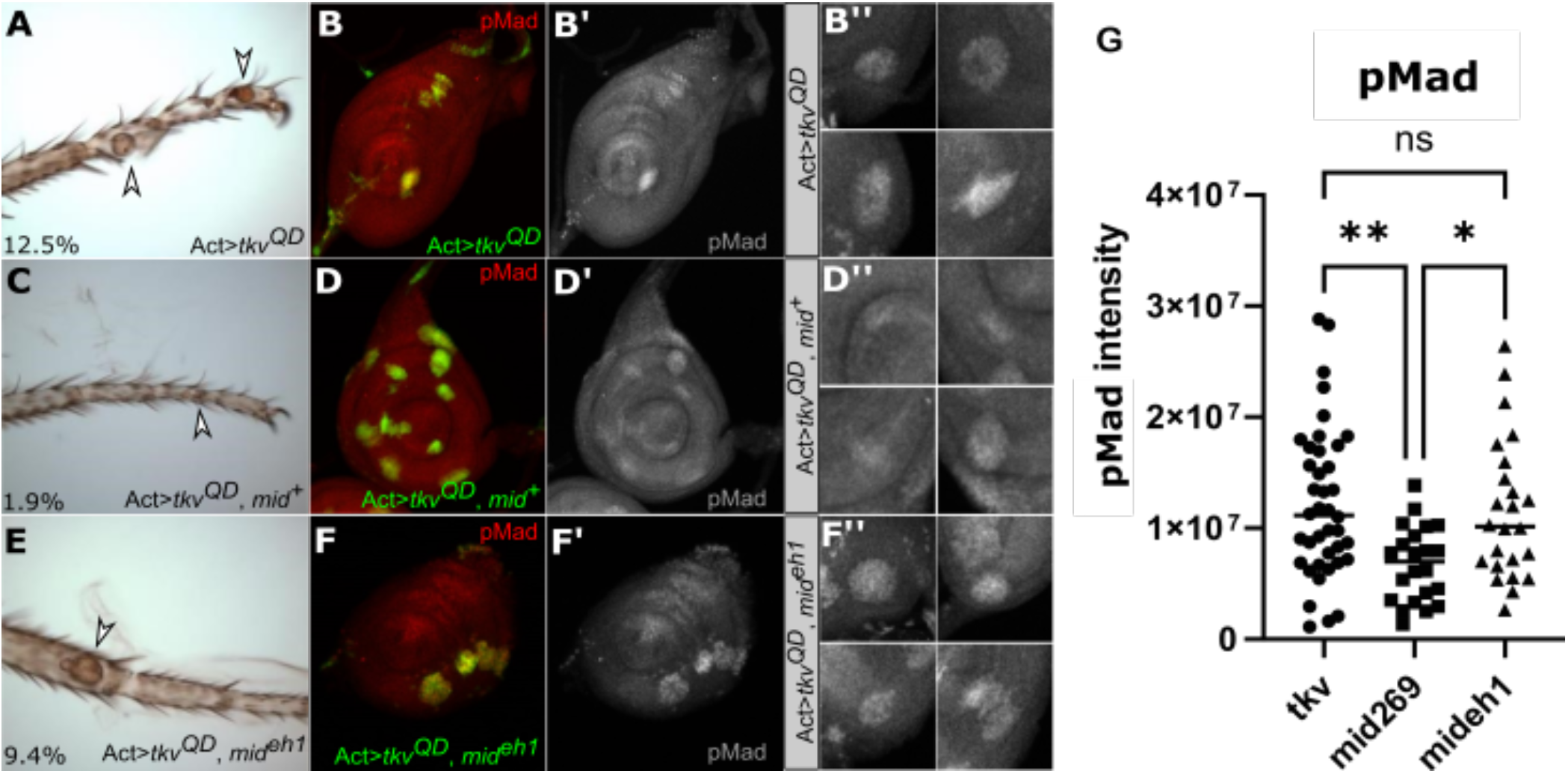
The eh1 domain is involved in Dpp repression. (A) AyGal4 gain-of-function clones expressing UAS-*tkv^QD^* result in outgrowths and dorsal transformation (arrowhead). 12.5% of legs had ectopic outgrowths under these conditions (n=181). (C) When UAS-*tkv^QD^* was co-expressed with UAS-*mid*^+^-Flag, the outgrowths were less severe (arrowhead) indicating suppression of the *tkv* gain-of-function phenotype. The effect was also less frequent with only 1.9% of legs having ectopic outgrowths (n=309). (E) Clones co-expressing UAS-*tkv^QD^* and UAS-*mid^eh1^*-Flag had deformities (arrowhead) similar to clones expressing UAS-*tkv^QD^* alone, where 9.4% of legs had ectopic outgrowths (n=106). pMad staining (red, greyscale single channel) was upregulated in clones (green) expressing UAS-*tkv^QD^* (B, B’, B”) and clones co-expressing UAS-*tkv^QD^* and UAS-*mid^eh1^*-Flag (green) (F, F’, F”). When UAS-*mid*^+^-Flag was expressed in clones (green) along with UAS-*tkv^QD^*, pMad staining was less elevated (D, D’, D”). (G) The difference in pMad levels staining between UAS-*tkv^QD^* clones and UAS-*tkv^QD^*, UAS-*mid^eh1^*-Flag was not significant. However, clones expressing UAS-*tkv^QD^*, UAS-*mid*^+^-Flag had significantly lower pMad compared to the other two conditions. Statistical analysis used the Tukey’s multiple comparisons test, with bars representing the mean and significance indicated, * (P value ≤ 0.05) and ** (P value ≤ 0.01).

In addition to the different patterning defects induced in adult legs, we also saw changes in the pMad accumulation in leg imaginal discs. Clones in third-instar discs expressing gain-of-function UAS-*tkv^QD^* have increased pMad staining (Fig. 4B,B’). Similar levels of pMad were detected in clones co-expressing UAS-*tkv^QD^* and UAS-*mid^eh1^*-Flag (Fig. 4F,F’). In comparison, clones expressing the UAS-*mid*^+^-Flag and UAS-*tkv^QD^* had lower levels of pMad staining relative to the other two genotypes (Fig. 4G). We note that the suppression of pMad by Flag-tagged Mid is less pronounced than by untagged UAS-*mid* (Svendsen et al., 2019) and this is consistent with our observation that all Flag-tagged Mid phenotypes are weaker compared to untagged Mid (Formaz-Preston et al., 2012; Svendsen et al., 2019). However, the UAS-*mid*^+^-Flag is able to rescue *mid* mutants, and the UAS-*mid*^+^-Flag and UAS-*mid^eh1^*-Flag transgenes are well matched for expression levels, with UAS-*mid^eh1^*-Flag expressed approximately 2-fold higher than UAS-*mid*^+^-Flag (Formaz-Preston et al., 2012). Together these results suggest that the eh1 domain is implicated in both of Mid’s repressive roles: direct repression of Dpp-target genes and interference with the Dpp-signaling cascade.

## Discussion

In this study, we investigated how the ventral selector gene *mid* antagonizes Dpp signaling. Specifically, we showed that *mid*, which antagonizes Dpp signaling and pMad accumulation, represses the Dpp-activated gene *Dad* but has no effect on the regulation of the Dpp-repressed gene *brk*. Furthermore, *mid* antagonizes dorsal fate by directly repressing *Dad* and by inhibiting Dpp signaling pMad accumulation via the eh1 domain.

We showed previously that Mid localizes to several Dad enhancers, including the *Dad13* enhancer, and represses a *Dad* enhancer trap in an eh1-dependent manner (Svendsen et al., 2019; unpublished data). Here we show that *mid* regulates Dad13 expression through a TBE site. Ventral Dad13 expression is increased and expanded in *mid* loss-of-function, while ectopic expression of *mid* blocks Dad13. Mutating the TBE in the Dad13 enhancer fragment increased Dad13 expression levels and expanded the ventral domain of expression compared to controls. Furthermore, the TBE mutation rendered the construct insensitive to *mid*^+^ gain-of-function, demonstrating that *Dad* is repressed by Mid through the TBE. Additionally, the dorsal expression of Dad13^TBE^ was increased compared to Dad13, suggesting that the TBE binding sequence binds other factors, perhaps inhibiting a dorsal repressor from binding or allowing the binding of activating factors, thereby affecting the dorsal expression.

Furthermore, we demonstrated that Mid suppresses pMad accumulation in a manner that is dependent on the eh1 domain. Mid mutants in which eh1 is compromised are unable to suppress Dpp gain-of-function effects. When co-expressed with Dpp-signaling overexpression in imaginal discs, clones of UAS-*mid^eh1^*-Flag maintain strong pMad staining and adult cuticles display outgrowths and deformities that resemble the effects of Dpp-signaling overexpression alone. Conversely, clones activated for Dpp signaling that express wildtype *mid* (UAS-*mid*^+^-Flag) have decreased pMad levels in imaginal discs, while adult legs have fewer and milder defects. This demonstrates that in addition to blocking Dpp-target genes such as *Dad, mid* represses genes that act to increase pMad levels and Dpp signalling. What these repressed target genes may be remains a subject for further study.

Our results showing that *brk* expression was not affected by *mid* were surprising to us because *brk* is a sensitive readout of Dpp signaling and because *mid* alters pMad levels. We showed that *mid* loss-of-function clones that increase pMad levels do not affect *brk* expression, even though *brk* is regulated by Dpp-signaling in ventral *mid*-expressing cells. It is possible that *brk’s* insensitivity to changes in *mid* is due to the negative feedback by *Dad* on Dpp-signaling in *mid* mutant clones. Though activated by Dpp, *Dad* is an inhibitory Smad (iSmad) that negatively regulates Dpp signaling (Tsuneizumi et al., 1997). Therefore, the induction of *Dad* creates an intracellular autoregulatory negative feedback loop which antagonizes Dpp. Thus, in *mid* loss-of-function, the increase in pMad may have little effect on the expression of *brk* because of the corresponding increased expression of inhibitory *Dad*. Additionally, *mid* expression was unable to suppress the repression of *brk* by Dpp gain-of-function, despite *mid* dramatically decreasing pMad accumulation in this genetic background (Svendsen et al., 2019). This persistence of *brk* repression may indicate that it is sensitive to even low levels of Dpp activation.

The lack of interaction between *mid* and *brk* in this study is consistent with previous work where it was reported that the expression of the ventral genes *H15* and *wg*, and the dorsal genes *omb* and *dpp*, were normal in *brk* mutant leg discs, indicating that *brk* does not participate in the formation of the DV axis (Estella and Mann, 2008). Instead, *brk* functions to form the proximo-distal (PD) axis by antagonizing the Wg target genes *Distalless (Dll)* and *dachshund* (*dac*), helping to establish their expression in the PD axis. This leads to a model in which Dpp induces the PD and DV leg axes through two separate modes: Dpp inhibits *brk* repression of Wg targets to establish the PD axis and antagonizes Wg targets in ventral development through pMad/Med/Shn mediated repression (Estella and Mann, 2008). Thus, *mid* and *brk* play parallel roles repressing Dpp-target genes in complementary regions of the ventral domain of the leg imaginal disc.

## Materials and Methods

### Fly stocks and constructs

All flies were maintained on standard media containing cornmeal, agar, yeast, glucose, and water (Deliu et al., 2017) and housed at between 18°C-25°C. Stocks UAS-*tkv^QD^*, *brk-lacZ*, and *Dad*-lacZ were procured from Bloomington Indiana Stock Center. *brk*-lacZ (BM315) was a gift from Dr. Konrad Basler (Müller et al., 2003). H15^X4^ mid^1a5^, UAS-*mid^V5^*, (Svendsen et al., 2009), UAS-*mid2.12* (Buescher et al., 2004), UAS-*mid^eh1^*-Flag and UAS-*mid*^+^-Flag constructs (Formaz-Preston et al., 2012) were previously generated. In this study, *mid*^+^ may be in reference to either UAS-*mid^V5^* or UAS-*mid2.12*. Dad13 constructs (Dad13, Dad13^TBE^, Dad13^SBE^, and Dad13^SBE+TBE^) were generated for this study with the use of the pGL3basic-hsp70-dad13 (331) plasmid gifted by Dr. Giorgos Pyrowolakis and mutations were made with an adapted splice protocol (Warrens et al., 1997) into a placZ-2.attB vector. The Dad13nRFP strain was a gift from Dr. Doug Allan.

### Loss-of-function and gain-of-function genetic mosaics

Heat shocking larvae 48-72hr after egg laying activates heat shock inducible hs-FLP, which is implemented in both gain-of-function and loss-of-function clonal experiments. In gain-of-function experiments, the combination of an AyGal4 construct with a UAS-linked gene allows for generation of ectopic clones which are labeled with GFP (Ito et al., 1997). Inducing the hs-FLP in loss-of-function experiments catalyzes mitotic recombination between FRT sites of homologous chromosomes (Xu and Rubin, 1993). This gives rise to two daughter cells homozygous for different genotypes, mutant and wildtype. The consequent clones have wildtype cells are marked with GFP, whereas loss-of-function cells lack of GFP.

### Reporter constructs, immunohistochemistry and imaging

The expression of *Dad*-lacZ (Tsuneizumi et al., 1997), *brk*-lacZ (Müller et al., 2003), and Dad13 constructs were detected with using mouse-anti-β-galactosidase (1:1000, Promega) and rabbit-anti-β-galactosidase (1:1000, Jackson ImmunoResearch Laboratories). *Brk* was also detected with rat anti-Brk (1:100, a gift from Gines Morata, Universidad Autonoma de Madrid). Dad13 was visualized by RFP expression. pMad was detected with the rabbit-anti-pSmad1/5/9 antibody (1:200, Cell Signaling Technology). Secondary antibodies used were Alexa-fluor 546 and 488 against rabbit or mouse (1:500, Molecular Probes). Imaginal discs were imaged on a Zeiss LSM 700 confocal microscope using the 20x objective with ZEN Black edition software. All compared genotypes were imaged at the same acquisition settings to maintain intensities. Adult tissues were visualized on a Leica MPS60 compound microscope with a 10x or 20x objective and QCapture_x64 software. Z-stack CZI files generated on the Zeiss LSM 700 confocal microscope were processed and analyzed with ImageJ software and saved as JPEG files. Some files were then further analyzed to determine fluorescence intensity in according to published methods (McCloy et al., 2014). Images were imported into INKScape to generate figures for this report. Graphs were generated and statistical analysis performed in GraphPad Prism 9 using ANOVA with Tukey’s multiple comparisons test.

## Acknowledgements

We thank Doug Allen (University of British Columbia), Konrad Basler (University of Zurich) and Indiana Stock Center for the Drosophila strains used in this study. Thank you to Gines Morata, (Universidad Autonoma de Madrid) for the *brk* antibody. Thank you to Giorgos Pyrowolakis (University of Freiburg) for the plasmid containing Dad13. We thank BestGene Inc. (Chino Hills, CA) for the transgene injection services.

## Competing interests

The authors declare no competing or financial interests.

## Funding

This work was supported by a National Sciences and Engineering Research Council of Canada (RGPIN/04828-2017 to WJB). LAP was funded by an Alberta Children’s Hospital Research Institute Award. LKK and MLA were supported by National Sciences and Engineering Research Council of Canada summer awards.

---

**Supplemental Figure 1.**
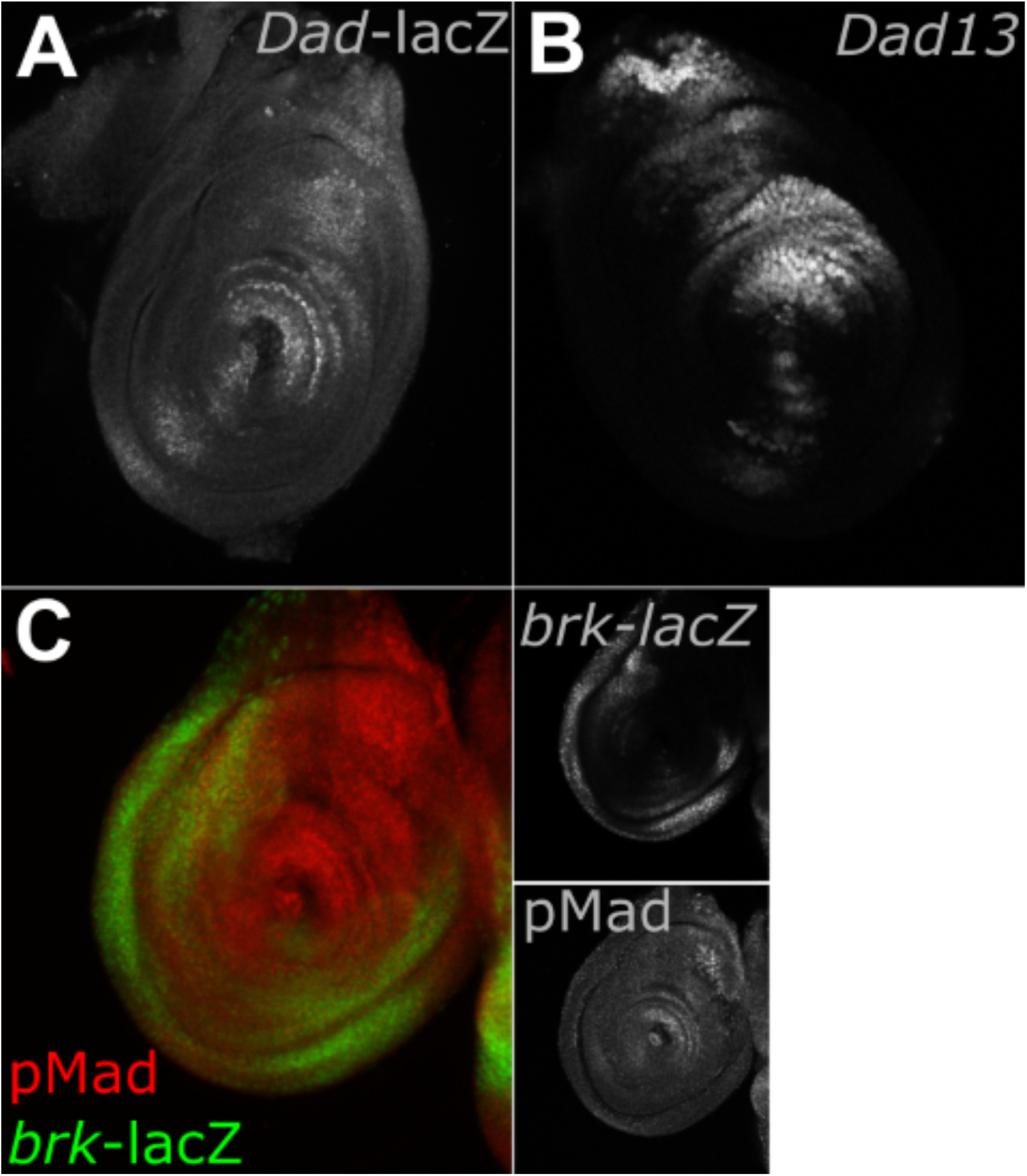
Expression patterns of *Dad, Dad13* and *brk*. (A-B) Both *Dad*-lacZ (A) and Dad13 (B) are expressed in the leg imaginal disc, with stronger expression in the dorsal domain and weaker expression the ventral domain. Dad13 (B) ventral expression is weaker than *Dad*-lacZ (A). (C) pMad and *brk* form a border in the leg imaginal disc with *brk* expression (green, top inset) ventral/lateral and pMad broadly expressed in the dorsal domain and narrowly expressed in the ventral domain (red, bottom inset).

**Supplemental Figure 2.**
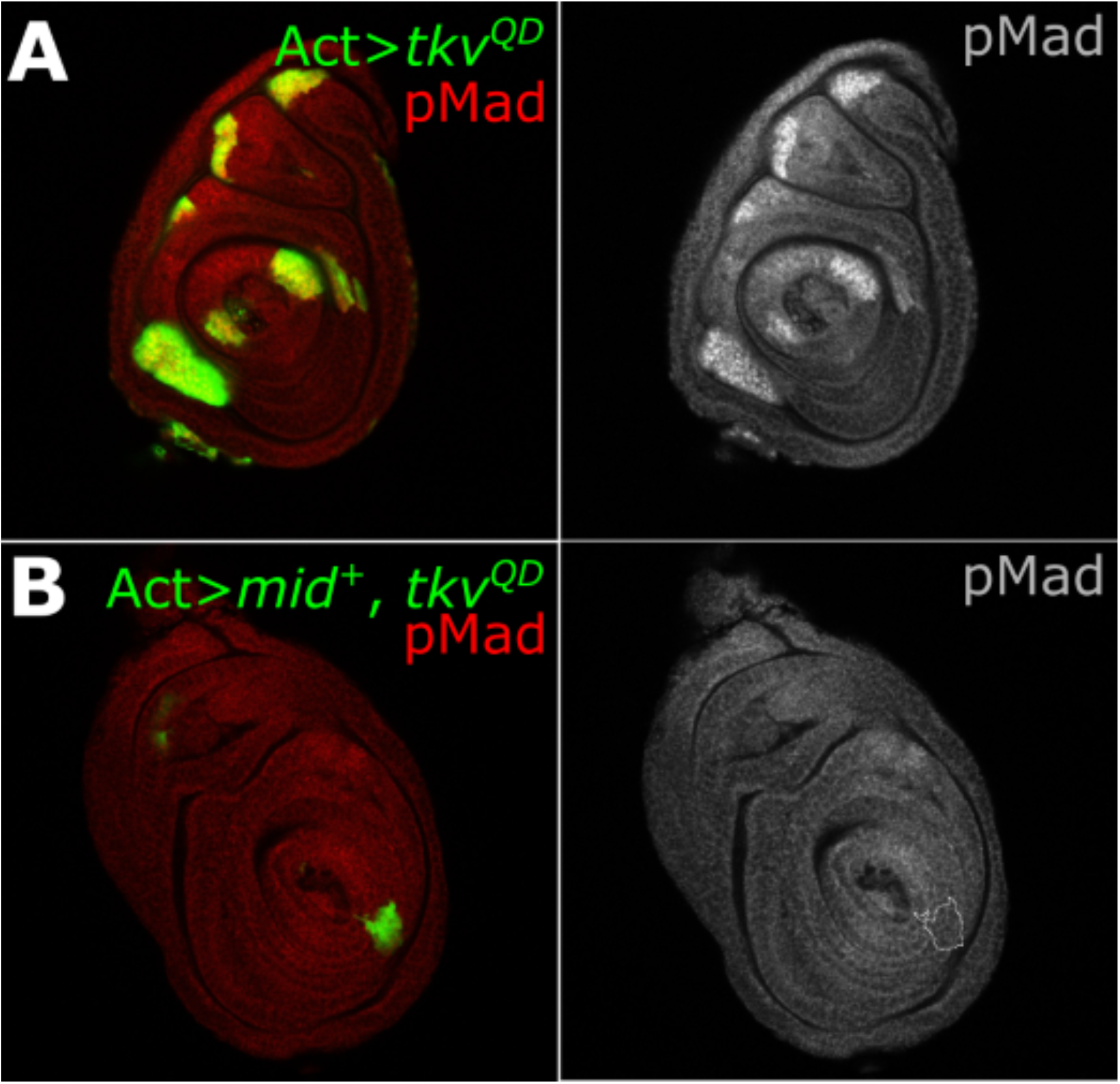
pMad supressed in clones co-expressing *mid* and Dpp signaling. (A) AyGal4 gain-of-function clones expressing UAS-*tkv^QD^* (green) results in outgrowths with increased pMad staining (red, single channel). (B) When AyGal4 clones are generated which express UAS-*mid*^+^ and UAS-*tkv^QD^* (green) the pMad staining is suppressed (red, single channel).

**Supplemental Figure 3.**
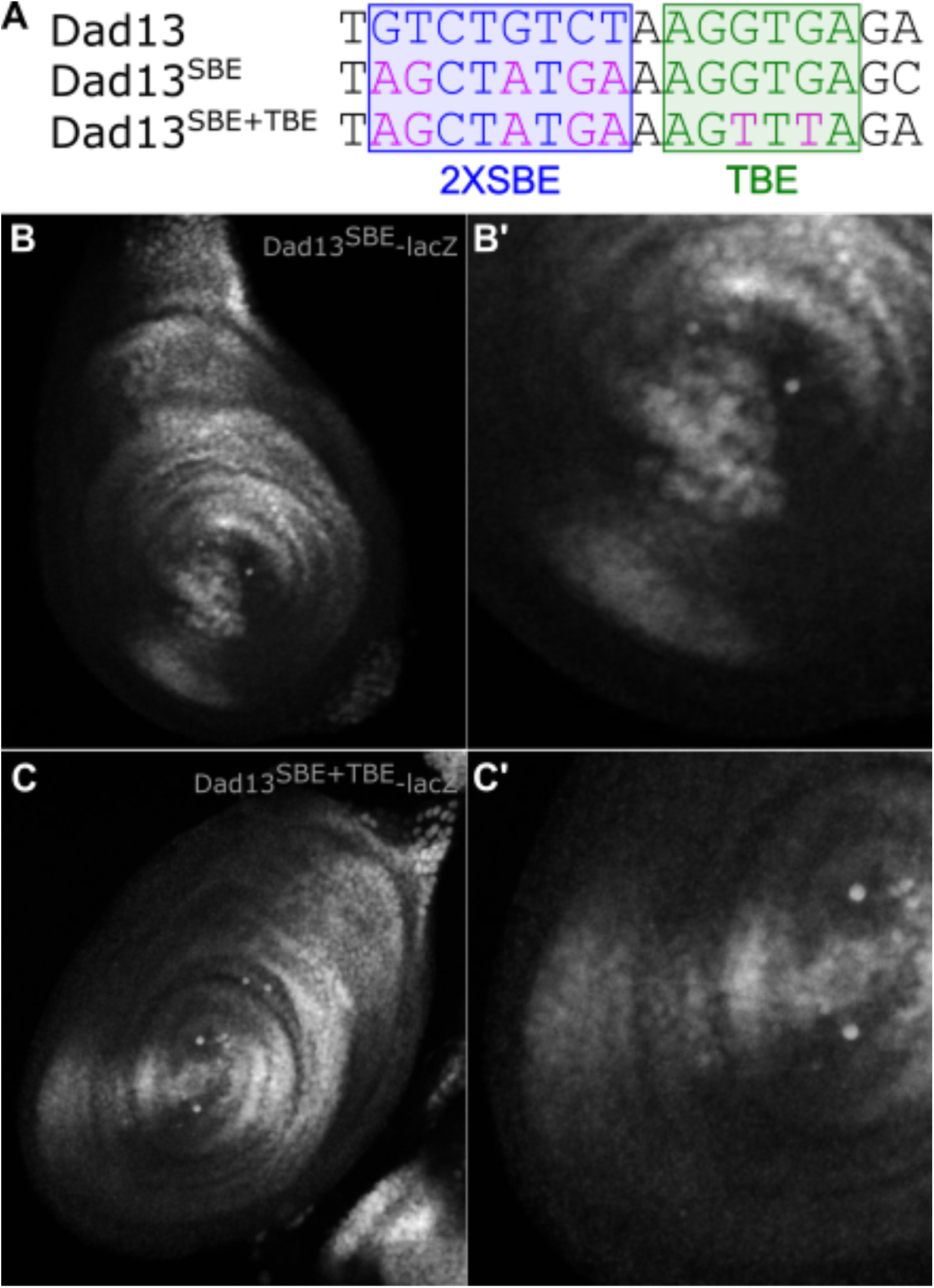
Expression patterns of other Dad13 constructs. (A) Part of the Dad13 enhancer fragment sequence, showing the 2X SBE sequence (blue) and the TBE sequence (green). Two G to T substitutions in the TBE were generated for the Dad13^TBE^ construct (fuchsia). Five nucleic acids were mutated (fuchsia) within the two SBE to generate the Dad13^SBE^ construct. A construct was also created that contained both the SBE and TBE mutations, termed Dad13^SBE+TBE^. (B) Dad13^SBE^ expression is similar to Dad13 expression, as detected by lacZ with strong staining in the dorsal domain and weak staining in the ventral domain. (B’) Magnified image of ventral expression of Dad13^SBE^-lacZ. (C) The double mutant construct, Dad13^SBE+TBE^, has an expression pattern similar to Dad13^TBE^ alone, with a wider and more intense expression in the ventral domain. Similar to Dad13^TBE^, Dad13^SBE+TBE^ dorsal expression is stronger than Dad13. (C’) Magnified image of ventral expression of Dad13^SBE+TBE^-lacZ.

**Supplemental Figure 4.**
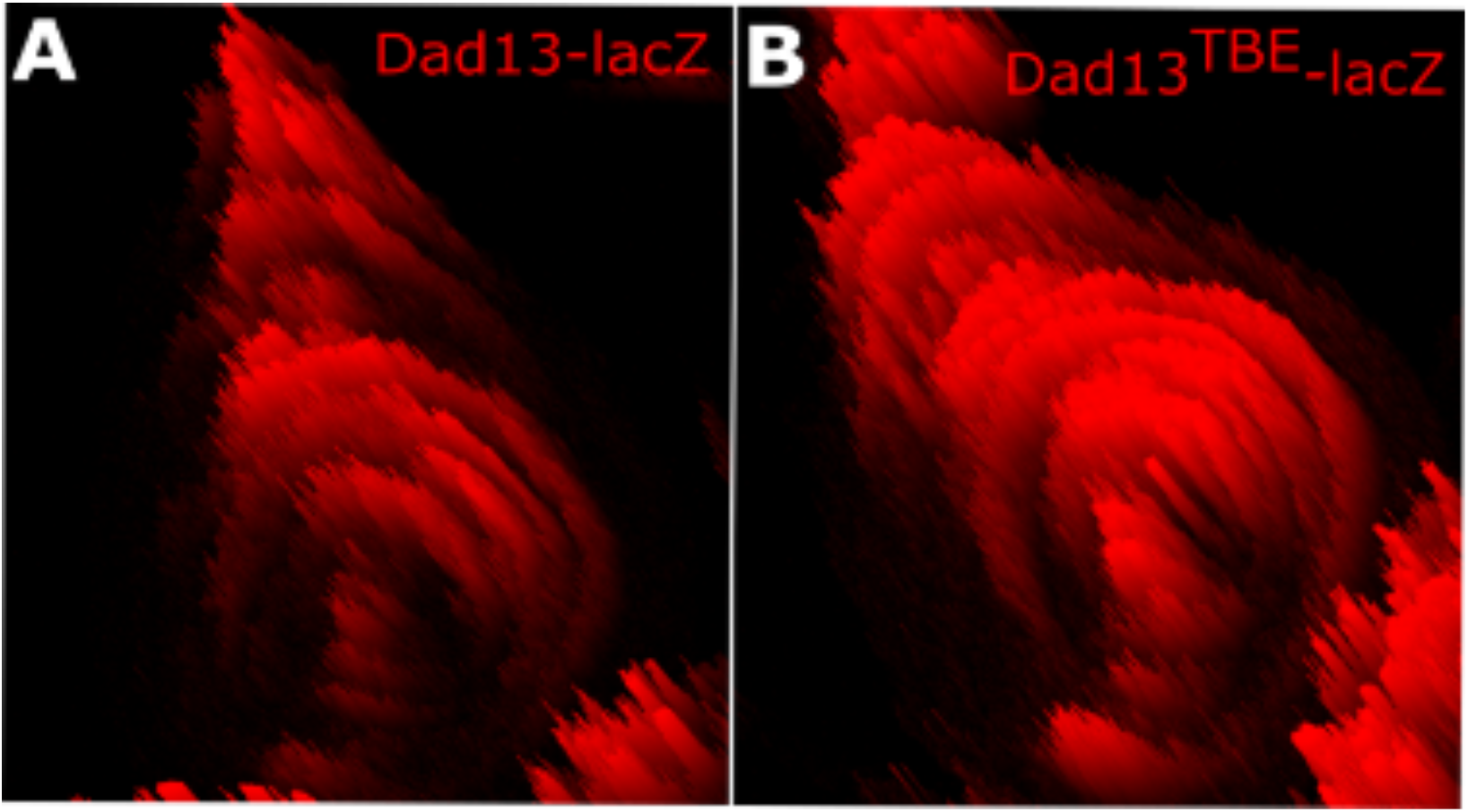
Dad13 and Dad13^TBE^ expression intensity. The 2.5D processing tool in ZEN was used to visualize intensity values of the lacZ staining for both Dad13 and Dad13^TBE^. (A-B) the expression of Dad13^TBE^ (B) is more intense in both the ventral and dorsal domain compared to Dad13 (A).

## Notes

### Competing Interest Statement

The authors have declared no competing interest.

